# Robustness of Transposable Element regulation but no genomic shock observed in interspecific *Arabidopsis* hybrids

**DOI:** 10.1101/258467

**Authors:** Ulrike Göbel, Agustin Arce, Fei He, Alain Rico, Gregor Schmitz, Juliette de Meaux

## Abstract

The merging of two divergent genomes in a hybrid is believed to trigger a “genomic shock”, disrupting gene regulation and transposable element (TE) silencing. Here, we tested this expectation by comparing the pattern of expression of transposable elements in their native and hybrid genomic context. For this, we sequenced the transcriptome of the *Arabidopsis thaliana* genotype Col-0, the *A. lyrata* genotype MN47 and their F1 hybrid. Contrary to expectations, we observe that the level of TE expression in the hybrid is strongly correlated to levels in the parental species. We detect that at most 1.1% of expressed transposable elements belonging to two specific subfamilies change their expression level upon hybridization. Most of these changes, however, are of small magnitude. We observe that the few hybrid-specific modifications in TE expression are more likely to occur when TE insertions are close to genes. In addition, changes in epigenetic histone marks H3K9me2 and H3K27me3 following hybridization do not coincide with TEs with changed expression. Finally, we further examined TE expression in parents and hybrids exposed to severe dehydration stress. Despite the major reorganization of gene and TE expression by stress, we observe that hybridization does not lead to increased disorganization of TE expression in the hybrid. We conclude that TE expression is globally robust to hybridization and that the term “genomic shock” is no longer appropriate to describe the anticipated consequences of merging divergent genomes in a hybrid.

## Introduction

Interspecific hybridization is an important factor in plant evolution: While fertile allopolyploid hybrids may become the founders of new species (Soltis & Soltis 2009), the merging of two divergent genomes in a hybrid can also cause mild outbreeding depression or even result in complete incompatibility (Todesco et al., 2016, Bomblies & Weigel 2007). At the molecular level, hybridization can lead to a genome-wide misregulation of the transcriptome and epigenome (Lafon–Placette & Köhler, 2015). The mobilization of up-regulated transposable elements (TE) can further result in chromosomal rearrangements. Such phenomena, first described by Barbara McClintock in maize (McClintock 1984) and termed “genomic shock”, are thought to provide a post-zygotic barrier against gene flow between species.

Yet, a recent study of the homoploid hybrid of *Arabidopsis thaliana* and *Arabidopsis lyrata* did not report a major disruption of the transcriptome and epigenome. Moderate differences in expression of protein-coding genes were reported and the parental pattern of the histone mark H3K27me3 appeared to be maintained (Zhu et al. 2017). These species, however, differ in both genomic structure and TE content. A recent increase in TE number has been reported for the *A. lyrata* genome (Hu et al. 2011). *A. thaliana* TEs are concentrated in pericentromeric regions, rarely venturing in gene rich chromosome arms. By contrast, in *A. lyrata* TEs account for most of the 40% larger genome and are broadly distributed, occurring in closer vicinity to expressed genes (Hu et al. 2011). The relatively low frequency of insertion polymorphisms within species revealed evolutionary tensions on insertions in gene rich regions, where permanent TE silencing can have deleterious consequences on the expression of neighboring genes (Lockton and Gaut, 2010, Hollister and Gaut 2009, Hollister et al., 2011). Finally, several studies have observed that *A. lyrata* TEs tended to be more highly expressed than *A. thaliana* alleles in hybrids (He et al. 2012). In view of these differences in number, distribution, and regulation of transposable elements, we hypothesized that for hybrids between these species, a genomic shock is more likely to occur at the level of TE expression. To test this hypothesis, we quantified the impact of parental vs. hybrid genomic background on TE regulation.

We studied transposable elements in the *A. thaliana* (Col-0) *x A. lyrata* (MN47) hybrid and its respective parental lines, using RNA-Seq expression data and ChIP-seq data of the histone marks H3K27me3 and H3K9me2. It is difficult to establish orthology relationships for TEs, because many of them duplicated after the species separation (Hu et al. 2011, de la Chaux et al., 2012). Thus our analysis focuses on trans-acting effects experienced by TEs of each genome after hybridization. Contrary to our primary expectations, we find TE silencing to be largely unaffected in the hybrid. No systematic relationship could be found between a change of TE expression in the hybrid and the gain or loss of histone marks. We further exposed interspecific hybrids and their parents to severe dehydration stress and confirm that TE regulation remains robust to hybridization in stress conditions.

## Materials and Methods

### Plant materials and growth conditions

Seeds of *A. thaliana* accession Col-0 were obtained from the Arabidopsis Biological Resource Center (ABRC, USA). *Arabidopsis lyrata* ssp. *lyrata* genotype MN47 was obtained from the lab of D. Weigel (Max Planck Institute for Development, Tübingen, Germany). Parental lines were crossed by pollinating emasculated *A. thaliana* flowers with *A. lyrata* pollen, as described in de Meaux et al. (2006). Reciprocal crosses using *A. thaliana* as pollen donor were not successful.

### RNA sampling and sequencing in standard and stress conditions

Seeds were stratified for 3 days at 4°C, germinated on soil and grown for 4 weeks in a growth chamber at 20°C under 14 h light/ 16°C 10 h dark under dim light (100 mmol sec-1 m-2). A dehydration treatment was applied following Seki et al. (2002). Plants were removed from the soil and dehydrated on paper for 0h and 3h, in the same growth chamber. The aerial part of the plant was sampled, flash-frozen in liquid nitrogen, and RNA was extracted as previously described (He et al. 2016). Four biological replicates were collected in one-week intervals. Two micrograms of total RNA were used for library preparation following the TruSeq^®^ Illumina RNA Sample Preparation v2 Guide. This includes poly-A+ RNA selection and the use of random primers for reverse-transcription. Sequencing was performed on Illumina HiSeq2000 following the manufacturer’s protocols and paired-end 100 bp long reads were obtained. 15-20 million paired reads for the parents and 30-40 million reads for the hybrid were produced (Suppl. Table 1). RNA-seq reads were filtered and trimmed for quality control as in He et al. 2016 and mapped on the hybrid genome, a concatenation of the *A. thaliana* Col-0 reference genome (TAIR10, www.arabidopsis.org) and the *A. lyrata* MN47 reference genome (Araly1, Nordberg et al. 2014)), using STAR with default parameters (Dobin et al. 2013).

### TE and gene read counting

The gene annotations TAIR10 (Col-0, Berardini et al. 2015), Alyrata_107_v1.0 (MN47, Hu et al. 2011) and TE annotations for *A. lyrata* and *A. thaliana* (Pietzenuk et al. 2016) were merged to form the genome annotation file used in this analysis. Aligned reads were filtered as follows: for the MN47 genome, only the main scaffolds 1-8 were considered. Read or fragment alignments shorter than 20 or longer than 700 base pairs (bp) were discarded. A minimum alignment score of E33 was required in order to accept an RNA-seq match. We allowed multiple RNA-seq read matches on TEs if they were entirely within a single (super)family, following Pietzenuk et al. 2016, and Jin et al. 2015. A read was assigned to a (super)family F on genome G if its primary match was in F on G, and any secondary matches on G were in F, too. In order to assign a match to a (super)family F, a minimum overlap of 50 bp to a member of F was required. Secondary matches on the alternative genome were allowed but not counted. Each read thus either contributed a single count to a (super)family or was discarded as multiply matching. If a read matched to TE_1_. TE_n_ of family F, the read was defined to contribute 1/n counts to each of these TEsU!Reads from a region of overlapping a TE and a gene were not discarded, but counted with the TE. These summed fractional per-element counts were the final output from the counting procedure. They were either used directly as counts per individual TE or the counts of the members of (super)families were summed up to yield aggregated per-(super)family counts. We implemented the counting procedure in the R programming language. After the counting step, annotated TEs without any read count were excluded from the analysis. Genes were counted with the same procedure as the TEs.

### Read count normalization

Read counts from genes and TEs were simultaneously normalized. Each count was first multiplied by a factor of 1000/L, where L is the length in base pairs of the respective annotated element (gene, individual TE, or entire TE (super)family). This removes the dependence of the read count on element length or (super)family size. The total length of a TE (super)family was defined to equal the summed length of all (super)family members considered, discounting any regions of overlap with other annotated elements.

### Differential expression analysis

The Generalized Linear Models (GLM) support of the R package edgeR (Robinson et al. 2010, McCarthy et al. 2012) was used to test the significance of differences in length-normalized counts between the hybrid and parental backgrounds. P-values were adjusted with the FDR correction (Benjamini & Hochberg 1995). For visualization in scatterplots and for the stringent definition of differential TE expression, length-normalized counts were converted to Counts per Million (CPM) and scaled (Robinson & Oshlack 2010) using the calcNormFactors function of the R package edgeR. This transformation mirrors what edgeR does internally when computing differential expression. It removes global count biases due to library size differences, easing the interpretation of the scatterplots and allowing the set-up of a sample-independent cutoff on the absolute expression value at 0 and 10 CPM.

### Analysis of distance distribution of TEs to neighboring genes

Distances of differential and non-differential TEs to the nearest gene or gene 5’-end were binned into classes 0-1 kb, 1-2 kb, 2-3 kb, and ≥ 3 kb. (TEs overlapping a gene were not considered.) A pseudocount of 1 was added to each bin count. A saturated log-linear model of the counts was computed using the R glm() function (family=”poisson”), using sum-to-zero contrasts. The p-value for distance d in Supplemental Table 1 is the Pr(>|z|) value of the interaction coefficient between the factors distance category and significance of hybrid- dependent TE expression.

### Random expectations of superfamily frequencies

To evaluate the enrichment of TEs changing their expression in hybrid background within each superfamily, we computed a random expectation. For this, we used the set of TEs showing differential expression in the hybrid background as a reference set and performed 10,000 random re-samplings of the same number of TEs from all TEs with evidence of expression. To account for the skewed distribution of these TEs in the vicinity of non-TE genes, both the reference set and the set of all TEs were binned by distance to the closest gene 5’-end. For each bin of the reference sample, the same number of TEs as contained in the bin was randomly selected from the corresponding distance bin of all TEs. The union of the re-sampled bins then formed one re-sampled set. The width of the bins increased with increasing distance: up to a distance of 1 kb, the bin width was 200 base pairs (bp), between 1 and 5 kb it was 500 bp, and beyond 5 kb it was 1000 bp, to avoid having bins with sparse numbers of TEs. A median, 5% and 95% quantile of each superfamily frequency was extracted from the 10,000 random samples and compared to the observed frequency of TEs with hybrid-dependent expression.

### Chromatin immune-precipitation experiments

Plants were germinated and grown on germination medium containing ½ Murashige and Skoog salts, 3% sucrose, and 0.8% agar. Seeds were stratified for 5 days at 4°C, and then transferred to a growth chamber under long-day conditions (14 hours cool white light supplemented with red light at 20°C, and 10 hours dark at 18°C). One week after germination, plants were transferred to soil and grown in the same chamber. Finally, leaves larger than 0.5 cm were sampled from 3 weeks-old Col-0, and 6-weeks-old MN47 and F1 hybrids for chromatin immunoprecipitation (ChIP).

ChIP experiments were performed with the MAGnify™ Chromatin Immunoprecipitation System (49-2024, Thermo Fisher Scientific) according to the manufacturer’s instructions, with the following modifications. Plant material was fixed in 20ml Crosslinking Buffer (0.4M Sucrose, 10mM Tris-HCl, pH 8.0, 10mM MgCl2, 1% formaldehyde) by applying vacuum twice for 15 minutes. Nuclei and chromatin were purified as in He et al. (2012). Chromatin was sonicated with a BioRuptor device (Diagenode, Liège, Belgium) for 30 times 30s at high setting with 30 s intermittent cooling in ice-water. A DNA fragment size of 300 to 600 bp was controlled by running an aliquot of de-crosslinked and purified DNA on 1.5% agarose gels. The following antibodies were used in immunoprecipitation: anti-rat IgG (R9255, Sigma;St. Louis, MO, USA), as well as anti-H3K27me3, anti-H3K9me2 and anti-H3 (07-449, 07-441 and 07-690, respectively, Millipore; Temecula, CA, USA). For each sample, ChIP-DNA from 8 experiments was pooled and purified by Qiagen MinElute Reaction Cleanup Kit. Finally, we obtained ChIP-DNA samples from Col-0, MN47 and Col-0xMN47 with H3K27me3 and H3K9me2, and Col-0 and Col-0xMN47 with H3.

### ChIP-seq library preparation and sequencing

DNA was sheared with Ion Shear reagents and the Ion Xpress Fragment Library Kit (Thermo Fisher Scientific, Waltham, Massachusetts) was used for subsequent library preparation. After ligation of the respective Ion Xpress barcode adapters, samples were amplified by 17 or 23 PCR cycles. Amplified libraries were quantified on Bioanalyzer DNA High-Sensitivity Chips (Agilent Technologies, Santa Clara, USA) and diluted to 9 pM. Emulsion PCR was performed on the Ion OneTouch System, followed by enrichment for template positive Ion Sphere Particles. Sequencing sample was loaded on an Ion Proton chip and sequenced on the Ion proton sequencer (Thermo Fisher Scientific) according to the manufacturer’s instructions.

### ChIP-seq read processing, mapping and peak detection

Raw data were pre-processed using the TORRENT SUITE version 4.0.1 (Thermo Fisher Scientific, Waltham, Massachusetts) to trim adapter sequences. Using the FASTX-Toolkit (http://hannonlab.cshl.edu/fastx_toolkit/), reads were then trimmed from the 3´ end to remove low quality bases (phred < 15). Reads shorter than 30 bases and reads of poor overall quality (more than half of the bases with phred < 20) were discarded. The mapping of the reads was performed with bowtie2 (Langmead and Salzberg, 2012), with the end-to-end parameter, without allowing mismatches in the seed (-N 0), and allowing for up to two alignments per read (-k 2); although then only the best alignment was kept. All samples were mapped to a hybrid reference genome obtained by concatenating the genomes of the parent as described above for RNAseq read mapping. Cross-hybridization between genomes was negligible. A summary of ChIP seq read data is given in Suppl. Table 2.

Quality assessment of the assays was performed using phantompeaktools (Kharchenko et al., 2008; Landt et al., 2012) to calculate NSC and RSC metrics (normalized and relative strand coefficients, respectively), and the bioconductor ChIPQC package (Caroll et al., 2014) to calculate the square sum of deviation. NSC was low but RSC metrics were well within the recommended range (Landt et al. 2012). Mapped reads were filtered with samtools (Li et al., 2009) for a mapping quality of 3 (-q 3), which discards multi-mapping reads with two equally good mappings but retains those with a single best mapping location. Duplicate reads from the amplification step during library preparation were removed with the MarkDuplicates program from Picard Tools (http://broadinstitute.github.io/picard). ChIP-seq read counts for TEs and non-TE genes were counted following the same pipeline as for RNAseq reads. A first analysis of ChIP-seq count distribution confirmed that the distribution of marks recovered known chromatin domains but revealed 64 genomic regions giving aberrant ChIP signals (see Suppl. Methods). These 64 regions were removed from all analyses presented in the manuscript.

## Results

### Only a minor fraction of all expressed TEs is differentially expressed in the hybrid

Our annotation contains 10863 TEs on the *A. thaliana* Col-0 genome and 30868 elements on the *A. lyrata* MN47 genome. After excluding genome-specific TE families, 10805 and 29150 elements remained on the Col-0 and MN47 genomes, respectively. Of these, 1592 (15 %) of the *A. thaliana* TEs and 7045 (24 %) of the *A. lyrata* TEs showed evidence of expression in at least one replicate. We quantified TE expression in normalized Counts Per Million (CPM, see methods). TEs contributed but a minor fraction of all reads (on average, only 0.07% in *A. thaliana* samples and 0.24% in *A.lyrata* samples). This fraction remained comparable in the hybrid transcriptome (on average 0.06% and 0.30% of *A. thaliana* and *A. lyrata* reads, respectively in the hybrid). Only a small fraction (maximally 2%) of the expressed TEs of both genomes were significantly differentially expressed between parental and hybrid backgrounds, using different cutoffs to define significance (Table 1). Moreover, this percentage dropped below 0.5% when the median expression was required to be above 10 CPM in at least one of the two backgrounds (Table1). These observations show that TE transcription is not severely disturbed in the hybrid.

**Table 1:** Numbers and fractions of differentially expressed TEs. At two different false- discovery rates (FDR <= 0.01 or <=0.05) and two different thresholds on the normalized Counts Per Million (CPM) (all: >=0 CPM; >=10 CPM). Numbers are given as “number of up-regulations” + “number of down-regulations”. Fractions were computed relative to all TEs with evidence of expression in standard conditions, discounting members of genome-specific families.

| | FDR $\leq 0.01$ | | FDR $\leq 0.05$ | |
| --- | --- | --- | --- | --- |
| | all | $\geq 10$ CPM | all | $\geq 10$ CPM |
| **A. thaliana** | 3+2<br>(0.3%) | 1+1<br>(0.1%) | 5+5<br>(0.6%) | 1+2<br>(0.2%) |
| **A. lyrata** | 57+2<br>(0.8%) | 14+0<br>(0.2%) | 73+6<br>(1.1%) | 18+1<br>(0.3%) |

### Specific rather than global effects act on TE expression in the hybrid

In order to investigate the effect of hybridization at the levels of individual elements, families and superfamilies of TEs, we correlated expression of individual TEs and combined expression levels of TE families between parents and hybrids (Fig. 1). While TE transcription in the two genetic backgrounds correlated well at all levels, the correlation increased from the individual TEs to the family and superfamily levels. Spearman’s rank correlation of the median expression of replicates in the two backgrounds under control conditions was 0.73, 0.89, 0.97 on the three levels of individual TE, family, and superfamily, respectively in *A. thaliana*, and 0.65, 0.86, 0.91, respectively in *A. lyrata*. Including four replicates in the RNA-seq analysis was necessary to establish these relationships with increased confidence. Using subsets of only two replicates, for example, reduced the parent-hybrid correlation coefficient of TE expression to 0.53, 0.77, 0.87 on the three levels for *A. thaliana*, and to 0.51, 0.84, 0.89, respectively for *A. lyrata*.

**Figure 1:**
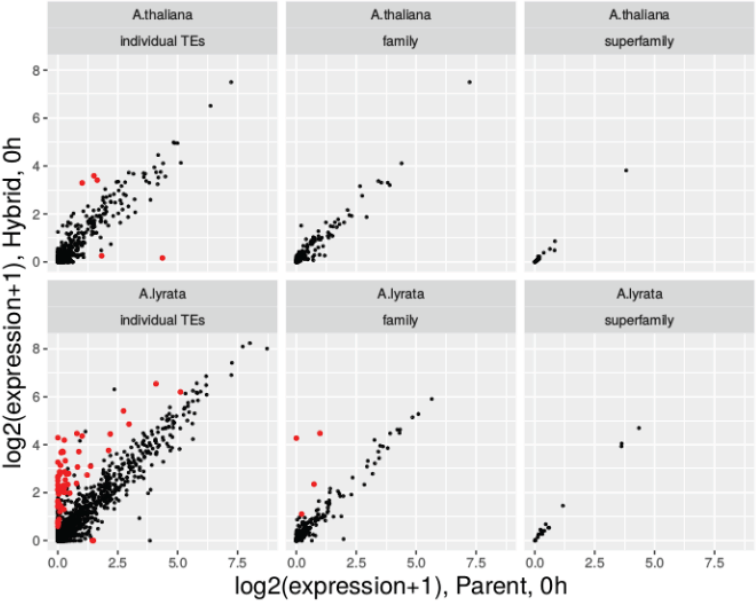
Correlation of TE expression levels, estimated as Median normalized counts per million over the replicate plants grown in standard conditions, in parent and hybrid for individual TEs, TE families, and superfamilies. TEs or (super)families with significant expression change are marked in red.

The small subset of TEs with significantly different expression between parent and hybrid was restricted to lower level units (individual TEs and families), indicating that the differences are case-specific rather than caused by a global disruption of TE silencing.

In *A. lyrata*, the 59 (0.8%) TEs that respond significantly to hybridization at FDR ≤ 0.01 were generally induced. In *A. thaliana*, the 5 (0.3%) affected elements were either induced or repressed (Fig. S1). SINEs dominated by Sadhu elements (Rangwala & Richards 2010) and Harbingers dominated by the ATIS112A family (Kapitonov & Jurka 2004) stood out as enriched among the responding *A. lyrata* TEs (Fig. 2). These superfamilies were significantly over- represented relative to the set of expressed TEs (≥ 1 RNA-seq count, permutation test, FDR < 0.05). The fact that only a small number of TE families showed enrichment for differential expressed elements indicates that hybridization does not have a global effect on TE expression.

**Figure 2:**
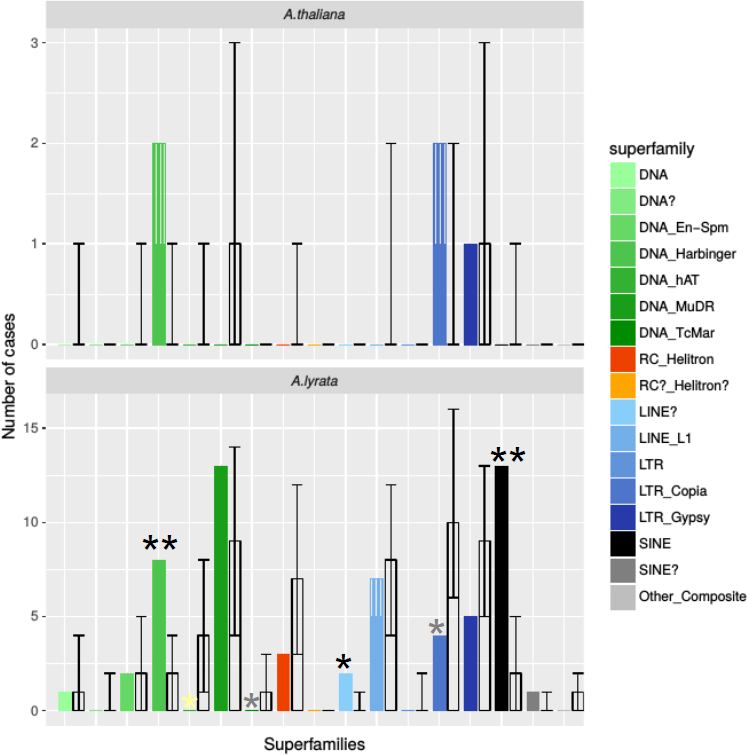
Observed and expected superfamily distribution of differentially expressed TEs (parent vs hybrid). Colored bars show the observed numbers of TEs showing hybrid- dependent expression in each TE superfamily. Each color corresponds to one family as indicated in the legend. Transparent bars indicate expected median count for random sampling within each superfamily. Error bars indicate the 5% and 95% quantile of the 10,000 random draws. Single black stars mark superfamilies whose observed count is significantly higher than expected from random expectations. Grey stars indicate depletion (maximally 5% of the universe have counts ≤ the observed count of the respective superfamily). Double stars indicate significant enrichment or depletion that is robust to FDR correction.

### Rare TE expression changes coincide with alterations in nearby gene expression

TEs can change expression of nearby genes by influencing the methylation patterns. We therefore asked whether hybrid-specific expression changes of TEs predict a corresponding change in neighboring genes.

Compared to TEs with expression not affected by hybridization, *A. lyrata* TEs expressed differentially in the hybrid were depleted in genome regions distant more than ≥ 3 kb from the closest gene 5’ ends (log-linear model, p ≤ 5 x 10-4, Table 2). In addition, counts of hybridization-sensitive TEs were enriched in the bins 1-2 kb upstream of gene 5’-ends (log- linear model, p ≤ 0.01, Table 2). This indicates a preferential location of hybridization-sensitive TEs in euchromatic, gene-rich regions in *A. lyrata*. In *A. thaliana*, there were too few differentially expressed TEs in the hybrid background to reliably compare the distance distributions.

**Table 2:** Distance distributions to nearby genes of *A. lyrata* TEs with differential expression in parent and hybrid under control conditions. The table combines distance information of TEs with a significant change in expression in the hybrid to either the nearest gene (any orientation) or the nearest 5’ of any gene. Numbers of significant and non-significant TEs are listed for each distance bin (FDR 0.01). In the lower part, p-values are given for the contribution of each distance bin to the difference between the distributions of significant and non-significant TEs (see Methods). Bins with significant depletion or enrichment in significant TEs are highlighted in blue and orange, respectively.

| distance to |  | closest gene |  |  |  | closest 5' end of a gene |  |  |  |
| --- | --- | --- | --- | --- | --- | --- | --- | --- | --- |
|  |  | <1 kb | 1-2 kb | 2-3 kb | ≥3 kb | <1 kb | 1-2 kb | 2-3 kb | ≥3 kb |
| Number of TEs | diff expr. P/H | 15 | 18 | 13 | 13 | 7 | 24 | 12 | 16 |
|  | not diff expr. | 1238 | 1675 | 913 | 3160 | 654 | 1455 | 1078 | 3799 |
| p (frequency in distance bin) |  | 0.239 | 0.539 | 0.061 | 4 e-4 | 0.554 | 0.010 | 0.517 | 1.5 e-4 |

We further analyzed if changed expression of TEs and their neighboring genes in a hybrid background were correlated. Hybrid-specific changes in transcription levels of non-TE genes have already been reported (Zhu et al. 2017). We also quantified non-TE gene expression levels in parents and hybrids. Note that in our experiment, the timing of sampling accounted for the developmental differences between the fast developer *A. thaliana* and both *A. lyrata* and the hybrid, which are late flowering. Markedly fewer genes were affected by hybridization in our experiment (234 in *A. thaliana* and 395 in *A. lyrata*), yet, we observed a significant overlap in genes reported to change expression in the hybrid background in Zhu et al. (2017, Hypergeometric test of overlap in genes changing their expression level between hybrid and parents, p<10-17 for all comparisons). In *A. lyrata* 153 TEs with detectable expression were within 2kb of a gene with hybridization-sensitive expression (FDR ≤ 0.01). Among these, 8 TEs displayed a change in expression in the hybrid background (p < 5 x 10-5, hypergeometric test). In the *A. thaliana* genome, the number of expressed TEs located in the vicinity of genes was too small to calculate correlations. This suggests that local chromatin alterations may alter the expression of both TE and gene in the vicinity. We conclude that hybrid-specific TE expression can coincide with a modification of the expression of nearby protein-coding genes, but this only affects a very small number of expressed loci.

### Histone marks are not modified in the hybrid background

Two epigenetic marks are associated with transcriptional silencing in plants: the plastic mark H3K27me3 and the permanent mark H3K9me2 (Sequeira–Mendes et al. 2014, Kim & Zilberman 2014). We profiled these two histone modifications by Chromatin Immuno-Precipitation followed by sequencing (ChIP-Seq) in hybrids and *A. thaliana* and *A. lyrata* parents. Although the ChIP-seq H3K27me3 yielded a comparatively lower number of reads, especially for Col-0 (Suppl. Table 2,5,6), the distribution of marks on TEs was in agreement with previous reports (Fig. 3A, Seymour et al. 2014) and captured well the chromatin domains that were defined in *A. thaliana* (Sequeira–Mendes et al. 2014, Suppl. Methods). In our data, H3K9me2 marks showed a high correlation of normalized read counts between parental and hybrid genomes, both genome-wide (calculated across 10kb genomic windows) and for individual TEs, TE families, and superfamilies. The genome-wide Spearman correlation coefficients of H3K9me2 read count along the genomes were 0.66 and 0.92 for *A. thaliana* or *A. lyrata*, respectively. Spearman’s rank correlation coefficients of H3K9me2 for individual TEs, TE-families, and superfamilies were 0.76, 0.94 and 0.96 for *A. thaliana* and 0.87, 0.97 and 0.99 for *A. lyrata* (Fig. 3B-C). While the high correlation values pointed to a largely additive inheritance of the mark in the hybrid on all levels of the annotation, the fact that correlation was lowest on the level of individual elements again underscores the role of element-specific responses in the hybrid.

**Figure 3:**
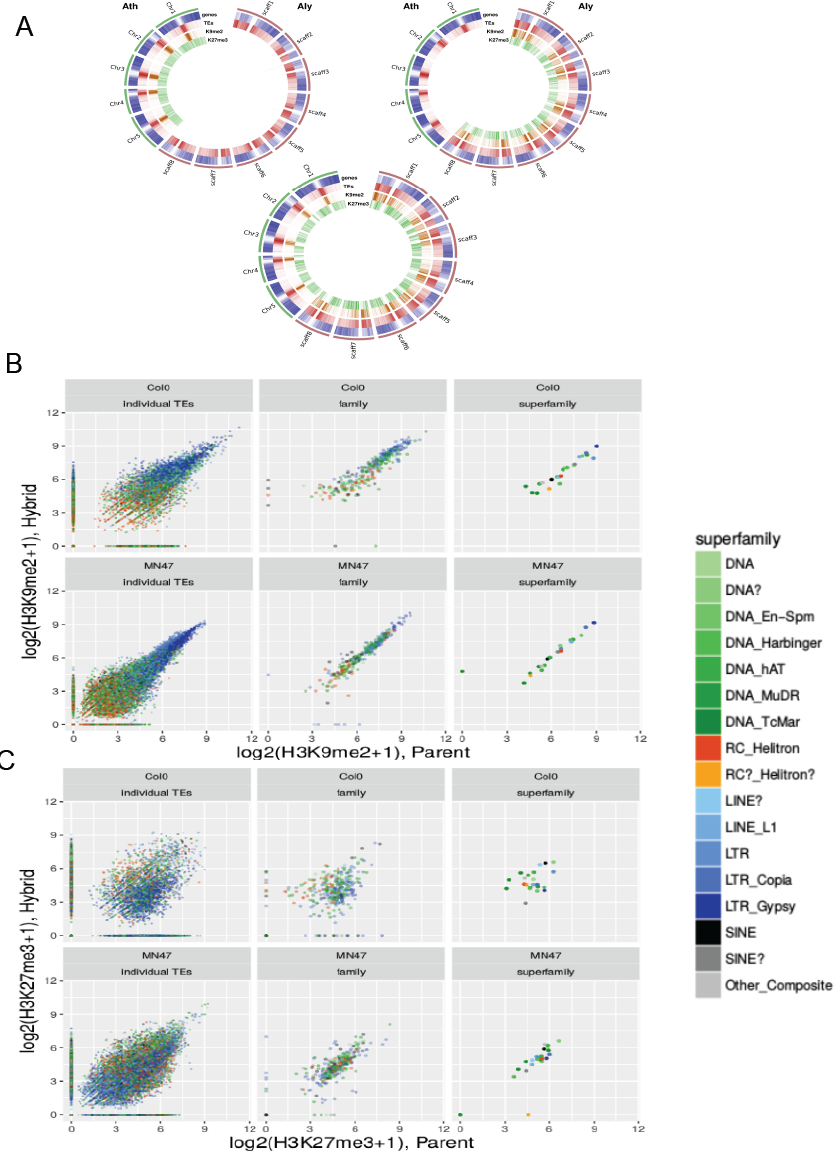
Correlation of histone mark levels on TEs in parent and hybrid. **(A)** Genome-wide distribution of H3K9me2 and H3K27me2 on the parental and hybrid genomes. The circular plots represent the hybrid genome used as reference, with the five *A. thaliana* chromosomes in green and the eight *A. lyrata* chromosomes in red. Inner tracks represent gene, TE, H3K9me2 and H3K27me3 densities in 500 kb bins. Top left: Col-0, Top right: MN47, Bottom: hybrid. **(B,C)** Scatterplots of ChIP-seq read counts. The color code indicates superfamily membership, using the palette of Figure 2. **(B)** H3K9me2 **(C)** H3K27me3.

While the function of H3K9me2 is to permanently silence TEs, H3K27me3 mainly serves to reversibly silence genes whose expression is restricted to specific conditions or developmental time windows (Gan et al. 2015). The distribution of H3K27me3 marks was strongly correlated genome-wide between parental and hybrid backgrounds (Spearman correlation coefficients: 0.63 and 0.86 for *A. thaliana* and *A. lyrata*, respectively, Fig. 3A). H3K27me3 is not a typical silencing mark of TEs (Sequeira–Mendes et al. 2014). This was most apparent in our data for the LTR superfamily, which tended to be associated with the highest level of H3K9me2 and the lowest level of H3K27me3 (Figure 3B, C). Nevertheless, the correlation of H3K27me3 occupancy in parent and hybrid remained high even for TEs, in *A. lyrata* (Spearman’s rank correlation 0.58, 0.78, 0.92 for individual TEs, families, and superfamilies). For the *A. thaliana*/hybrid comparison, the correlation dropped to 0.2, 0.27, 0.21 for individual TEs, families, and superfamilies, respectively (Fig. 3C). Note however that the Col-0 H3K27me3 ChIP-seq experiments yielded a comparatively lower number of reads
(Suppl. Table 2).

We tested then whether changes in histone mark occupancy could explain TE expression differences between parents and hybrids. TEs with a modified expression level in hybrid background generally showed similar H3K27me3 level in parent and hybrid (Fig. 4). When a departure in epigenetic levels was detected between parents and hybrids, the direction of the change in expression of the TE was not necessarily in the expected direction (Fig. 4). *A. lyrata* TEs with a modified expression level in hybrid background also generally showed similar H3K9me2 mark levels in parents and hybrids. We detected a concordant loss of H3K9me2 marks and increased expression in the hybrid for 6 *A. lyrata* TEs (Fig. 4). But there again, we observed for 2 TEs that the change in H3K9me2 occupancy did not reflect the change in expression. Altogether, in our data, a differential deposition of epigenetic marks in the hybrid is rare and generally does not coincide with hybrid-dependent changes in TE expression.

**Figure 4:**
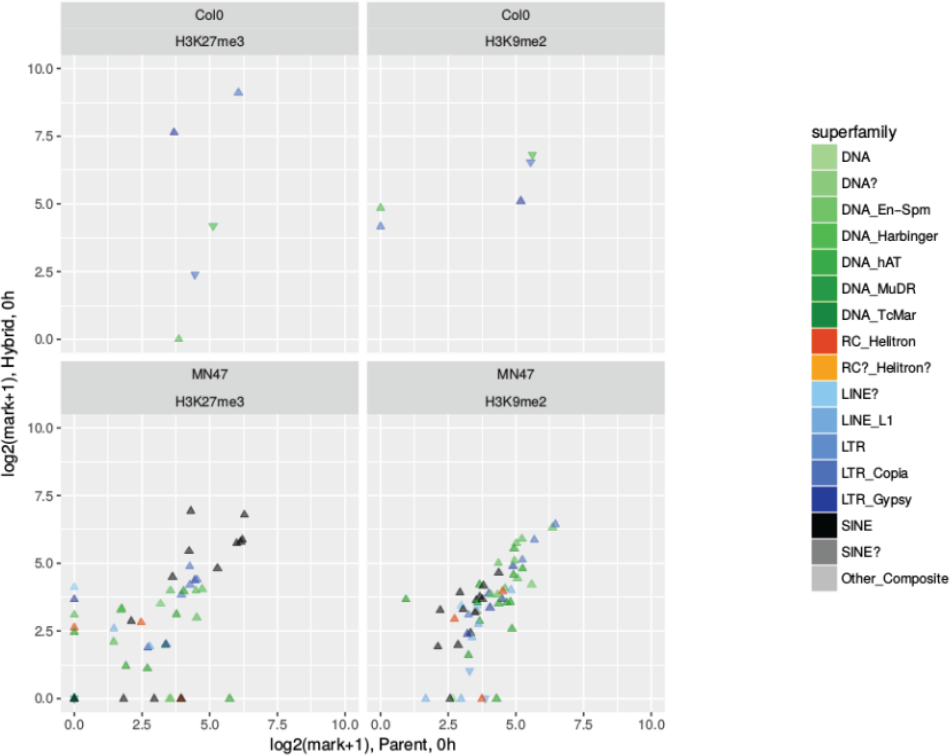
No change in histone mark occupancy is detected for differentially expressed TEs. Shown are scatterplots of each histone marks on each of the two genomes for TEs with significantly different expression in parent vs hybrid under standard conditions. Superfamily membership of the TEs is indicated using the palette of Figure 2. Symbol shapes indicate the direction of gene expression change in hybrid vs. parent (upward triangle: up, downward: down).

### Robustness of TE expression to hybridization is maintained under stress conditions

It is assumed that transposition activity is induced by stress conditions (McClintock 1984, Cavrak et al. 2014). Dehydration stress is associated with a vast transcriptome reprogramming (Matsui et al. 2008). We analyzed the dependence of transcript abundance upon interspecific hybridization for TEs and non-TE genes under strong drought stress conditions in the same way as described for control conditions. Having found that TEs with a hybrid specific expression change tend to be close to genes, and given the reprogramming of the gene transcriptome by the stress, we asked whether TE regulation was robust to hybridization even under stress conditions. In our data, 30-40% of the genes of both species changed expression after 3h of dehydration, with similar proportions in the parental and hybrid backgrounds. The fraction of stress responsive TEs was much smaller: 3-4% on the *A. thaliana* genome and 2-3% for *A. lyrata* (not shown). While the fraction of *A. thaliana* TEs with a significant expression difference between parent and hybrid is higher under dehydration stress than under control conditions, both the absolute numbers (4 up, 14 down) and the fraction (1.1% of all TEs with read count ≥1) remained small (Table 3). The fraction of responding *A. lyrata* TEs decreased slightly compared to control conditions, and again the numbers remained small (33 up, 18 down, fraction 0.7%). The percentage dropped below 0.5% if an expression level of ≥10 CPM in at least one of the backgrounds was required (Table 3). Like under control conditions, the correlation of TE expression levels between parents and hybrids increased at the family and superfamily levels. The Spearman’s rank correlation of the replicates’ median expression in the two backgrounds was (0.7, 0.81, 0.83) for *A. thaliana*, and (0.67, 0.91, 0.94) for *A. lyrata*, for individual, family and superfamily levels, respectively. Interestingly, we observed that more TEs were down-regulated in the hybrids compared to the parents under stress conditions than under standard conditions (Fig. 5A, Table 3). Only the Harbinger and LINE_L1 superfamiles were enriched among expressed *A. lyrata* TEs in stress conditions (Fig. 5B; permutation test using TEs with ≥ 1 RNA- seq count as reference, FDR < 0.05). This further corroborates the conclusion that in our two species, TE regulation is robust to hybridization even under stress conditions.

**Figure 5:**
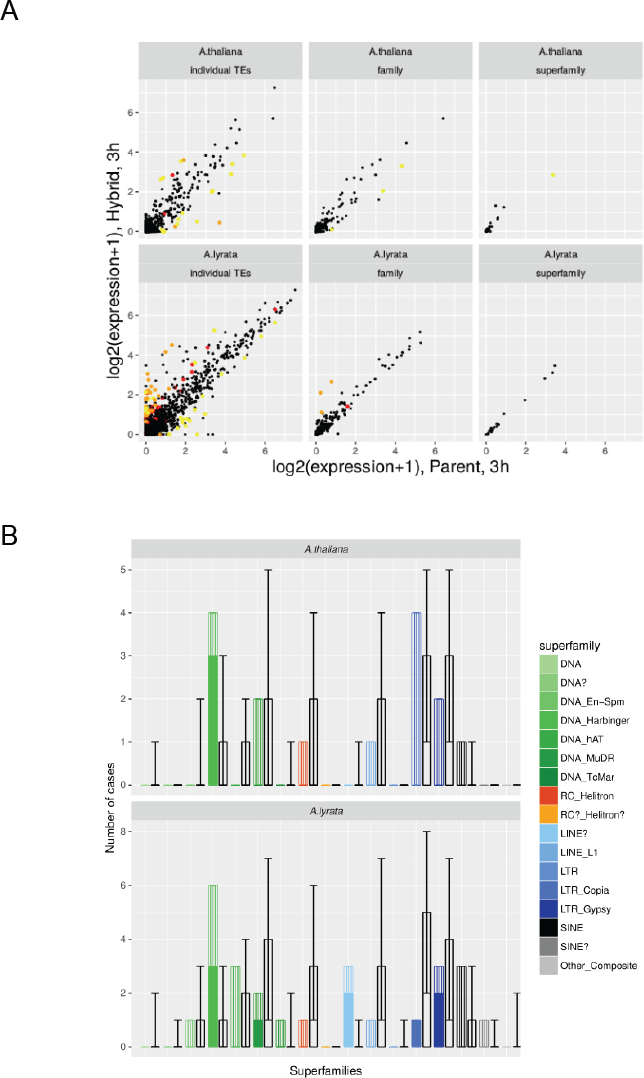
**(A) Correlation of TE expression levels in stress conditions**, estimated as Median normalized counts per million over the replicates, in parent and hybrid for individual TEs, TE families, and superfamilies. TEs or (super)families with significant expression change are marked in red (significant only under control conditions), orange (significant both under standard and stress conditions), or yellow (significant only in stress conditions). **(B) Observed and expected superfamily distribution of TEs whose expression is hybrid- dependent specifically in stress conditions**. Colored bars show the observed numbers of TEs showing hybrid-dependent expression in each TE superfamily. Each color corresponds to one family as indicated in the legend. Transparent bars indicate expected median count for random sampling within each superfamily. Error bars indicate the 5% and 95% quantile of the 10,000 random draws. Single black stars mark superfamilies whose observed count is significantly higher than expected from random expectations. Grey stars indicate depletion (maximally 5% of the universe have counts ≤ the observed count of the respective superfamily). Double stars indicate significant enrichment or depletion that is robust to FDR correction.

**Table 3:** Numbers and fractions of differentially expressed TEs in stress conditions (3h) and in response to stress (3h specific), At two different false-discovery rates (FDR <= 0.01 or <=0.05) and two different thresholds on the normalized Counts Per Million (CPM) (all: >=0 CPM; >=10 CPM). “Fractions were computed relative to all TEs with evidence of expression under either standard and/or stress conditions, discounting members of genome-specific families.

|  |  | FDR ≤ 0.01 |  | FDR ≤ 0.05 |  |
| --- | --- | --- | --- | --- | --- |
|  |  | all | ≥10 CPM | all | ≥10 CPM |
| A. thaliana | 3h | 4+14<br>(1.1%) | 2+4<br>(0.4%) | 7+26<br>(2.1%) | 2+5<br>(0.4%) |
|  | 3h specific | 3+12<br>(0.9%) | 2+3<br>(0.3%) | 4+23<br>(1.7%) | 2+4<br>(0.4%) |
| A. lyrata | 3h | 33+18<br>(0.7%) | 4+4<br>(0.1%) | 54+37<br>(1.3%) | 10+15<br>(0.4%) |
|  | 3h specific | 9+17<br>(0.4%) | 3+4<br>(0.1%) | 12+34<br>(0.7%) | 7+14<br>(0.3%) |

## Discussion

### No evidence for a genome shock upon hybridization of A. lyrata and A. thaliana

The term „genomic shock, was initially used to describe the breakage and large scale rearrangement of chromosomes in maize (McClintock 1946, McClintock 1984). This term is thus appropriate to describe genome instability following interspecific hybridization, i.e. unbalanced segregation of homoelogous chromosomes, chromosome loss or translocations (Chester et al. 2012, Spoelhof et al. 2017, Henry et al. 2014, Xiong et al. 2011). Over the years, the term “genome shock” has also been used to describe the anticipated alteration in genome activity following interspecific hybridization or allopolyploidization, even if in most cases the merging of divergent genome does not cause genome instability. In plants, DNA cytosine methylation (mC) and the histone mark H3K9me2 embed TEs in a chromatin context unfavorable for transcription (Kim & Zilberman 2014). The establishment and maintenance of these marks is not only inherited through DNA replication, it is also guided by small interfering RNAs (siRNAs), (see Kim and Zilberman 2014 for a review of plant TE silencing mechanisms). Because of their potential to exert genome-wide *trans* effects via base pair homology of the short 24nt siRNAs, the population of siRNAs expressed by a genome must be adapted to its gene content to ensure proper gene expression. Interspecific hybridization, therefore, brings together trans-regulators of TE expression that are not mutually adapted and could deeply disorganize gene regulation. Gene expression modifications following interspecific hybridization or allopolyploidization have been monitored across plant genera as diverse as *Gossypium, Senecio, Brassica, Tragopogon or Oryza (* e. g. Buggs et al 2009, Hegarty et al 2006, Yoo et al. 2013). Many studies have concluded that epigenome incompatibilities are important contributors to hybrid sterility (Senerchia et al. 2016, Wu et al. 2016, Wu et al. 2015, Ghani et al. 2014, Xu et al. 2014, Wang et al. 2014, Parisod et al. 2009). In *Aegilops* hybrids, for example, parental divergence in TE content was associated with the emergence of new TE copies in hybrids, while in tomato, transgressive expression of small RNAs was associated with the dysregulation of TEs (Senerchia et al. 2015, Shivaprasad et al. 2012).

Our analysis contradicts the conclusions of these previous reports. Comparing the TE expression and epigenetic profile of the *A. lyrata* and *A. thaliana* genomes in parental and hybrid context, we find no evidence for a major transcriptomic or epigenomic “shock” driven by inappropriate TE silencing and epigenetic mechanisms. Both TE transcription and the decoration of TEs by the silencing histone mark H3K9me2 are globally unaffected in the hybrid. We observe that only a handful of specific TEs show a change in expression in hybrids. In addition, the changes in TE expression are generally of small magnitude. We further show that the distribution of silencing epigenetic marks is strongly correlated in parents and hybrids, and its variation does not match the changes in TE expression. Finally, under severe dehydration stress, a large number of genes change their transcriptional activity (Matsui et al. 2008). This broad genome-wide remodeling of transcriptional activity, however, far from inducing a severed genome shock, tends to increase the robustness of TE expression patterns to hybridization. These findings thus suggest that stress does not increase the impact of hybridization, further supporting the idea that the control of TE elements is robust to genome merging. Although these observations partially contradict previous reports, they are not isolated. In recent years, an increasing number of studies, mostly conducted in the Brassicaceae family, has begun to report cases where hybridization does not massively alter gene regulation (Akama et al. 2014, He et al. 2016, Shi et al. 2015, Zhang et al. 2016, Zhu et al. 2017).

The contradiction between these studies and previous reports may have various explanations. First, it is striking to note that many of the studies that conclude on the absence of a global gene expression dysregulation were conducted in species with well-known genomes, where gene expression levels can be quantified with the best accuracy. They were not conducted in polyploids with large and imperfectly assembled genomes. Second, biological and experimental variance between replicates affects the strength of the correlation of expression in parental vs. hybrid backgrounds, and thus the apparent global impact of hybridization. Finally, it is necessary to contrast diverse sources of information to test whether hybridization has a broad impact on gene regulation. In the allotetraploid *A. suecica*, modification in siRNA levels was not associated with non-additive gene expression in newly formed allopolyploids, suggesting that changes in siRNA expression were not the result of epigenetic disorganization in the hybrid (Ha et al. 2009). Similarly, in *Arabidopsis* homoploid hybrids, we observed that hybrid-specific TE expression changes do not coincide with obvious epigenetic changes, a pattern also reported for protein-coding genes (Zhu et al. 2017). Taken together, the different molecular datasets used in this study show that TE expression and the distribution of epigenetic marks, which control gene expression, are not massively disorganized in hybrids.

### Trans- and cis Eacting differences between species can manifest in F1 hybrids

The comparative analysis of gene expression in hybrids and their parents reflects the variability of trans- and cis-regulatory mechanisms controlling gene expression in the two species (Wittkopp et al. 2004). After hybridization, changes in total expression will simply reflect the number and magnitude of *trans*-regulatory differences between species, whereas the specific *cis* Eregulatory variant of each transcript will result in allelic expression differences. These differences will manifest in hybrids on a restricted number of genes or elements, which does not imply that there is a major “genomic shock”.

Only *trans*-acting differences will have non-additive consequences on expression. In fungi and cotton allopolyploid, more than 50% of homeologous genes inherit the expression pattern of the parents (Cox et al. 2014, Yoo et al. 2013). In rice, allelic expression bias in hybrids between *Oryza sativa* and *O. japonica* also reflects mostly interspecific differences. In hybrids between the diploid *O. sativa* and the tetraploid *O. punctata*, only 16% of the genes showed an expression pattern different from that of the parents (Wu et al. 2016). The analysis of protein-coding gene expression in *A. thaliana* x *A.lyrata* hybrids reported a larger proportion of non-additive gene expression changes for the *A. thaliana* genome compared to the *A. lyrata* genome (Zhu et al. 2017). This pattern, like in our study, could not be associated with chromatin modifications induced by hybridization or any other genome-wide change in gene regulatory mechanisms (Zhu et al. 2017). However, the *A. thaliana* parental genotype Col-0 is missing an active allele of the major developmental regulator *FRIGIDA*. In fact, the *A. thaliana* allele of *FLC*, which is up- regulated by *FRIGIDA*, is one of the alleles most strongly impacted by hybridization (Suppl. Table 3, Zhu et al. 2017). Interspecific differences in *trans*-acting factor activity will therefore determine the extent to which gene expression changes upon hybridization. If F1 hybrids are fertile, these changes can segregate in subsequent F2 generations as in e.g. Shivaprasad et al. (2012).

Here, focusing on the *trans*-acting effect of the passage from a native parental background to a hybrid background on TE expression, we show that at most 1-2% of expressed TEs have expression that is differentially controlled in *trans*. Several elements indicate that these *trans*- acting effects may be associated with interspecific differences in mechanisms of gene regulation in euchromatic regions. When combined in a hybrid, these trans-acting differences modify the regulatory environment of a few TE sub-families. Indeed, the small proportion of TEs with differential expression in hybrids tend to be located in the vicinity of genes. Interspecific differences in the regulation of euchromatic regions are however likely to be minor, given the small number of elements affected.

Interspecific differences in gene regulation controlled in *cis* can also create a pattern of genome dominance in hybrids, with a majority of orthologous transcripts showing a transcription bias towards one parental genome (Cheng et al. 2016, Zhang et al. 2016, Yoo et al. 2013). For example, in *A. thaliana x A. lyrata* hybrids, a bias towards preferential expression of the *A. lyrata* allele has been reported by independent studies (He et al. 2012, He et al. 2016, Zhu et al. 2017). *Cis*-effects have not been examined in this study, because orthologous relationships cannot be established for most TEs, but for the subset of TEs that are located in orthologous positions in both species, *A. lyrata* TEs tend to be more highly expressed (He et al.2012).

### Incompatibilities can also have an oligogenic basis

It may seem at first surprising that TE silencing shows so little disruption in interspecific *A. thaliana x A. lyrata* hybrids, given the estimated 6M years of divergence and their vastly different TE content (Hu et al. 2011, Hohmann et al. 2015). The probability to observe specific disruptions in gene expression in a hybrid background, however, does not necessarily scale with evolutionary distance. The proportion of genes with hybrid-specific changes in expression is not greater for hybrids between the closely related sub-species *O. sativa* and *O. japonica* than between *O. sativa* and the more distant species *O. punctata* (Wu et al. 2016). A simple oligogenic combination of alleles, in fact, can cause what is known as a Dobzhansky-Muller incompatibility, even in the absence of large evolutionary distances between parents. Dobzhansky-Muller incompatibilities, which can cause major disruption at the phenotypic and, probably as a result at the transcriptome level, are not seldom within species, and were even detected within populations (Bomblies & Weigel 2010, Alcázar et al. 2010. Chae et al. 2014, Wright et al. 2013, Durand et al. 2012). We conclude that the term “shock” is no longer appropriate to describe the phenotypic consequences of the merging of divergent genomes in a hybrid. Our data rather supports a less dramatic scenario, where the merging of two genomes creates a new *trans*- acting background, in which oligogenic Dobzhansky-Muller incompatibilities can manifest. Without a global robustness of TE regulation to genome merging, hybridization and polyploidization would probably be much less important for plant diversification (Alix et al. Annals Bot. 2017).

## Supplementary Material

**Supplementary Methods:** Methods for quality control of ChIP seq data.

**Supplementary Figures 1:**
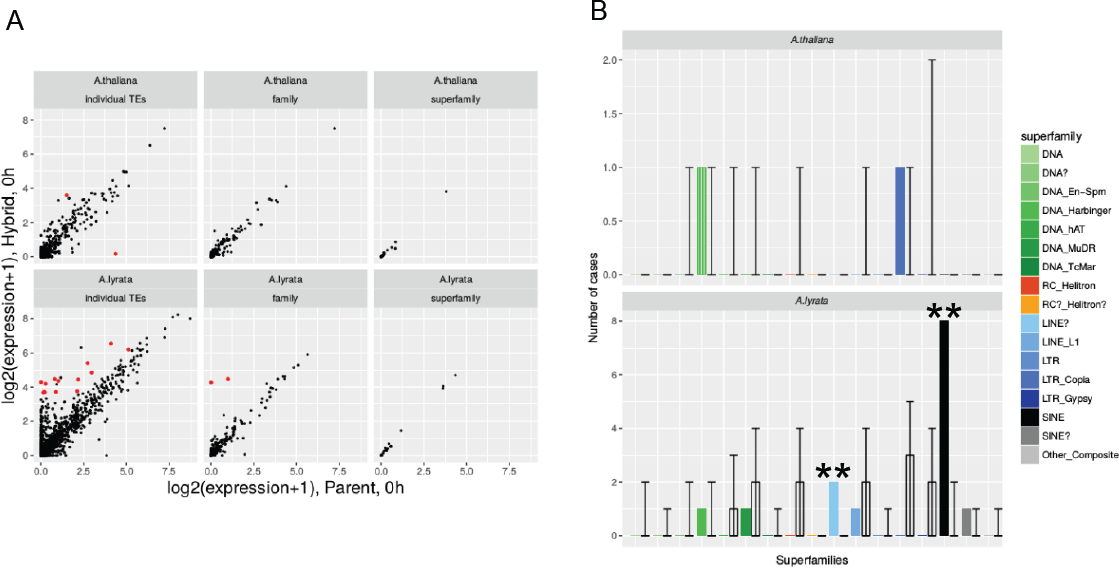
**(A) Correlation of TE expression levels**, estimated as Median normalized counts per million over the replicate plants grown in standard conditions, **in parent and hybrid** for individual TEs, TE families, and superfamilies expressed above a threshold of 10 CPM. TEs or (super)families with significant expression change are marked in red. (**B) Observed and expected superfamily distribution of differentially expressed TEs (parent vs hybrid).** Colored bars show the observed numbers of TEs showing hybrid-dependent expression above a threshold of 10 CPM in each TE superfamily. Each color corresponds to one family as indicated in the legend. Transparent bars indicate expected median count for random sampling within each superfamily. Error bars indicate the 5% and 95% quantile of the 10,000 random draws. Single black stars mark superfamilies whose observed count is significantly higher than expected from random expectations. Grey stars indicate depletion (maximally 5% of the universe have counts ≤ the observed count of the respective superfamily). Double stars indicate significant enrichment or depletion that is robust to FDR correction.

**Supplementary Figures 2:**
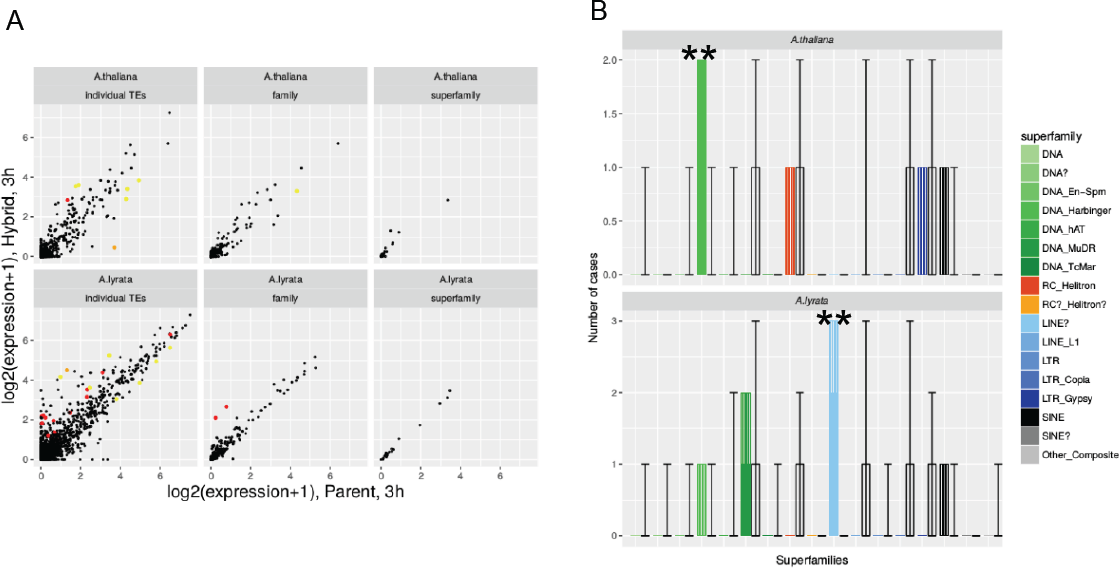
**(A) Correlation of TE expression levels in stress conditions**, estimated as Median normalized counts per million over the replicates, in parent and hybrid for individual TEs, TE families, and superfamilies expressed above a threshold of 10 CPM. TEs or (super)families with significant expression change are marked in red (significant only under control conditions), orange (significant both under standard and stress conditions), or yellow (significant only in stress conditions). **(B) Observed and expected superfamily distribution of TEs whose expression is hybrid-dependent specifically in stress conditions**. Colored bars show the observed numbers of TEs showing hybrid-dependent expression above a threshold of 10 CPM in each TE superfamily. Each color corresponds to one family as indicated in the legend. Transparent bars indicate expected median count for random sampling within each superfamily. Error bars indicate the 5% and 95% quantile of the 10,000 random draws. Single black stars mark superfamilies whose observed count is significantly higher than expected from random expectations. Grey stars indicate depletion (maximally 5% of the universe have counts ≤ the observed count of the respective superfamily). Double stars indicate significant enrichment or depletion that is robust to FDR correction.

### Supplementary tables

**Supplementary table 1:** Summary RNA-seq reads

**Supplementary table 2:** Summary ChIP-seq reads

**Supplementary table 3:** Median RNA-seq read counts for individual TEs and genes in parent and hybrid background in standard and stress conditions. Counts are normalized to 1 kb of element length. Raw and FDR-adjusted p-values for differential expression in the hybrid are given unless a TE or a gene had been excluded from the differential expression analysis by the low expression filtering.

**Supplementary table 4:** As Supplementary table 4, but only listing TEs and collapsing element counts by family

**Supplementary table 5:** ChIP-seq read counts for individual TEs and genes in parent and hybrid background. Counts are normalized to 1 kb of element length. Raw and FDR-adjusted p-values for differential expression in the hybrid are given unless a TE or a gene had been excluded from the differential expression analysis by the low expression filtering.

**Supplementary table 6:** As Supplementary table 5,but only listing TEs and collapsing element counts by family.

## Acknowledgements

We thank Franziska Turck (Max Planck Institute for Plant Breeding Research) for sharing her ChIP-seq protocol and experience. This research was supported by the Deutsche Forschung Gesellschaft (DFG) with grant ME2742/2-1 and ME2742/6-1 in the realm of the special study program SPP1529, as well as by the European Research Council with Grant 648617 “AdaptoSCOPE”. Raw data has been deposited in NCBI’s Gene Expression Omnibus (Edgar *et al.* 2002) and are accessible through GEO Series accession number (to be provided).

